## Supplementary Information for "Not getting in too deep: a practical deep learning approach to routine crystallisation image classification"

| | |
|---|---|
| Initial Learning Rate | $2e^{-4}$ |
| Learning Rate Factor | 0.5 |
| Batch Size | 16 |
| Optimizer | Adam optimiser |
| Loss Function | Cross-Entropy |
| Epochs | 100 |
| Horizontal/Vertical flipping | Yes |
| Zoom Range | 30% |
| Rotation Range | 30° |
| Width/Height Shifting | 5% |
| Re-scaling | 1/255 |

**Table S1:** Optimal training and image augmentation parameters used to train classifiers with each chosen network architecture.

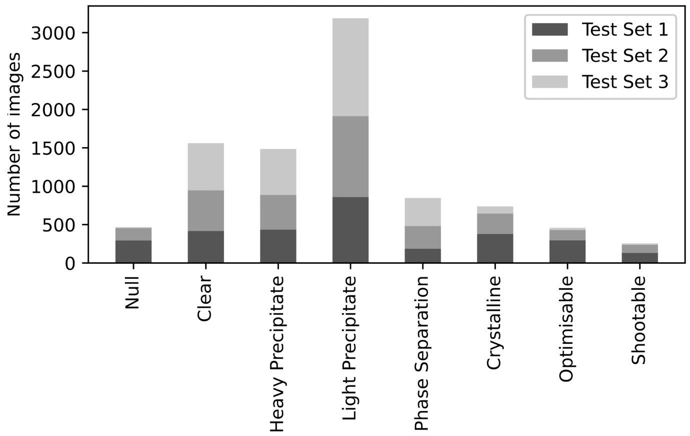


**Figure S1:** The number of images in each class for the three test data sets.

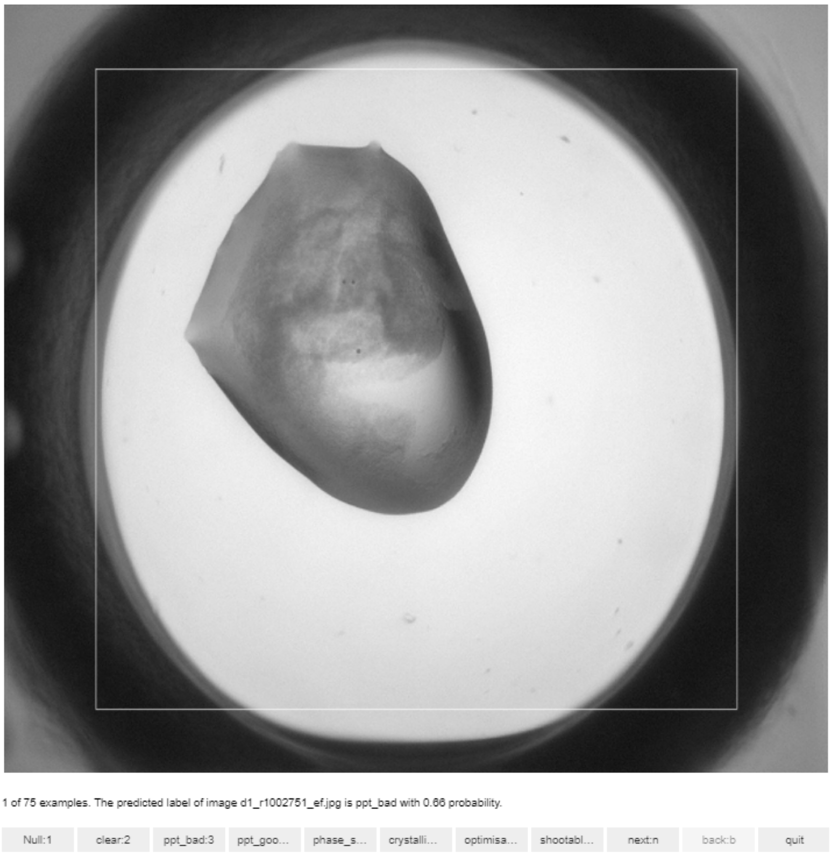


**Figure S2:** Screenshot showing the GUI used to check the results of labelling.

| Test Sets | Test 1 | | | | Test 2 | | | | Test 3 | | | |
|---|---|---|---|---|---|---|---|---|---|---|---|---|
| Classes | DenseNet121 | ResNet50 | InceptionV3 | Xception | DenseNet121 | ResNet50 | InceptionV3 | Xception | DenseNet121 | ResNet50 | InceptionV3 | Xception |
| Top-1 Accuracy (%) | 87.8 | 75.4 | 82.3 | 84.7 | 80.1 | 72.1 | 72.9 | 76.5 | 87.2 | 77.7 | 79.3 | 76.5 |
| Adjacent Accuracy (%) | 93.3 | 89.3 | 94.5 | 95.1 | 85.2 | 84.2 | 85.9 | 87.6 | 92.4 | 86.3 | 93.7 | 95.3 |
| Kappa | 0.86 | 0.71 | 0.79 | 0.82 | 0.75 | 0.66 | 0.67 | 0.71 | 0.83 | 0.69 | 0.73 | 0.77 |
| F1 | 0.89 | 0.74 | 0.83 | 0.85 | 0.81 | 0.71 | 0.73 | 0.77 | 0.88 | 0.79 | 0.79 | 0.83 |
| Precision | 0.89 | 0.81 | 0.86 | 0.86 | 0.85 | 0.78 | 0.79 | 0.81 | 0.89 | 0.84 | 0.85 | 0.86 |

**Table S2:** The test results of all classes for each architecture and test set.

| Test set 1 | null | clear | heavy-ppt | light-ppt | phase-sep | crystalline | optimisable | shootable |
|---|---|---|---|---|---|---|---|---|
| null | 77.6 | 10.2 | 1.7 | 2.7 | 3.1 | 2.4 | 1.7 | 0.7 |
| clear | 0.0 | 96.2 | 0.0 | 1.0 | 0.5 | 1.9 | 0.5 | 0.0 |
| heavy-ppt | 0.5 | 0.9 | 74.9 | 13.1 | 2.8 | 7.6 | 0.2 | 0.0 |
| light-ppt | 0.2 | 0.8 | 1.9 | 90.6 | 3.4 | 2.9 | 0.2 | 0.0 |
| phase-sep | 0.0 | 0.0 | 0.0 | 0.0 | 95.2 | 4.3 | 0.5 | 0.0 |
| crystalline | 0.0 | 0.0 | 0.3 | 0.3 | 2.4 | 95.3 | 1.6 | 0.3 |
| optimisable | 0.3 | 0.0 | 0.0 | 0.0 | 2.4 | 15.2 | 79.4 | 2.7 |
| shootable | 0.0 | 0.0 | 0.0 | 0.0 | 0.0 | 2.3 | 1.5 | 96.2 |

| Test set 2 | null | clear | heavy-ppt | light-ppt | phase-sep | crystalline | optimisable | shootable |
|---|---|---|---|---|---|---|---|---|
| null | 71.4 | 13.0 | 0.6 | 4.3 | 2.4 | 6.3 | 1.2 | 0.6 |
| clear | 0.2 | 89.8 | 0.2 | 0.2 | 0.6 | 8.1 | 0.9 | 0.0 |
| heavy-ppt | 1.1 | 0.2 | 58.2 | 13.5 | 2.1 | 23.5 | 0.9 | 0.4 |
| light-ppt | 0.0 | 0.9 | 1.0 | 82.1 | 6.3 | 8.4 | 1.1 | 0.1 |
| phase-sep | 0.0 | 0.0 | 1.7 | 0.0 | 93.2 | 4.1 | 0.0 | 0.0 |
| crystalline | 0.0 | 0.4 | 0.4 | 1.5 | 6.0 | 89.1 | 1.1 | 1.5 |
| optimisable | 0.0 | 0.0 | 5.9 | 4.4 | 8.1 | 20.7 | 52.6 | 8.1 |
| shootable | 0.0 | 0.0 | 0.9 | 0.0 | 1.9 | 5.6 | 0.9 | 90.7 |

| Test set 3 | null | clear | heavy-ppt | light-ppt | phase-sep | crystalline | optimisable | shootable |
|---|---|---|---|---|---|---|---|---|
| null | 66.7 | 0.0 | 0.0 | 0.0 | 16.7 | 16.7 | 0.0 | 0.0 |
| clear | 0.0 | 94.1 | 0.0 | 0.2 | 0.6 | 1.5 | 4.1 | 0.0 |
| heavy-ppt | 0.2 | 0.0 | 74.7 | 17.7 | 2.8 | 3.8 | 0.7 | 0.0 |
| light-ppt | 0.0 | 0.2 | 2.1 | 86.6 | 7.8 | 3.1 | 0.2 | 0.0 |
| phase-sep | 0.0 | 0.0 | 0.0 | 0.5 | 98.9 | 0.5 | 0.0 | 0.0 |
| crystalline | 0.0 | 0.0 | 1.1 | 5.5 | 0.0 | 93.4 | 0.0 | 0.0 |
| optimisable | 0.0 | 7.4 | 0.0 | 29.6 | 0.0 | 7.4 | 55.5 | 0.0 |
| shootable | 0.0 | 0.0 | 0.0 | 0.0 | 0.0 | 0.0 | 0.0 | 100.0 |

**Figure S3:** Confusion matrix showing the results for each of the three test sets obtained using the DenseNet121 classifier. Rows show class labels with predicted class in columns.

| Test set 1 | null | clear | heavy-ppt | light-ppt | phase-sep | crystalline | optimisable | shootable |
|---|---|---|---|---|---|---|---|---|
| null | 86.8 | 2.4 | 3.1 | 0.3 | 0.0 | 0.0 | 7.5 | 0.0 |
| clear | 13.9 | 85.2 | 0.0 | 0.2 | 0.0 | 0.0 | 0.7 | 0.0 |
| heavy-ppt | 0.7 | 3.0 | 67.4 | 26.4 | 1.1 | 0.5 | 0.9 | 0.0 |
| light-ppt | 2.8 | 3.5 | 2.4 | 87.7 | 2.2 | 0.2 | 1.2 | 0.0 |
| phase-sep | 1.1 | 3.2 | 0.0 | 2.1 | 86.1 | 0.5 | 7.0 | 0.0 |
| crystalline | 0.8 | 6.1 | 3.4 | 12.4 | 6.1 | 26.1 | 45.1 | 0.0 |
| optimisable | 0.0 | 0.3 | 0.0 | 0.0 | 0.0 | 0.7 | 99.0 | 0.0 |
| shootable | 0.8 | 0.8 | 0.0 | 2.3 | 0.0 | 0.0 | 58.0 | 38.2 |

| Test set 2 | null | clear | heavy-ppt | light-ppt | phase-sep | crystalline | optimisable | shootable |
|---|---|---|---|---|---|---|---|---|
| null | 87.0 | 3.1 | 3.7 | 1.2 | 2.5 | 0.6 | 1.9 | 0.0 |
| clear | 2.8 | 95.5 | 0.0 | 0.2 | 0.8 | 0.0 | 0.8 | 0.0 |
| heavy-ppt | 1.8 | 4.4 | 66.6 | 12.2 | 4.9 | 1.5 | 8.6 | 0.0 |
| light-ppt | 2.5 | 4.1 | 6.5 | 76.8 | 8.2 | 0.2 | 1.8 | 0.0 |
| phase-sep | 0.7 | 2.0 | 0.7 | 2.0 | 75.3 | 0.3 | 19.0 | 0.0 |
| crystalline | 3.0 | 5.2 | 13.1 | 15.0 | 13.5 | 12.4 | 37.8 | 0.0 |
| optimisable | 0.0 | 0.0 | 0.0 | 2.2 | 2.2 | 0.0 | 95.6 | 0.0 |
| shootable | 0.9 | 0.9 | 0.0 | 2.8 | 2.8 | 0.9 | 70.1 | 21.5 |

| Test set 3 | null | clear | heavy-ppt | light-ppt | phase-sep | crystalline | optimisable | shootable |
|---|---|---|---|---|---|---|---|---|
| null | 100.0 | 0.0 | 0.0 | 0.0 | 0.0 | 0.0 | 0.0 | 0.0 |
| clear | 2.4 | 97.4 | 0.0 | 0.0 | 0.0 | 0.2 | 0.0 | 0.0 |
| heavy-ppt | 1.7 | 2.5 | 64.2 | 25.8 | 0.3 | 0.0 | 5.5 | 0.0 |
| light-ppt | 3.8 | 4.2 | 3.4 | 86.0 | 1.3 | 0.8 | 1.3 | 0.0 |
| phase-sep | 11.3 | 4.4 | 0.0 | 9.6 | 54.7 | 1.1 | 19.0 | 0.0 |
| crystalline | 4.4 | 33.0 | 0.0 | 24.2 | 1.1 | 24.2 | 13.2 | 0.0 |
| optimisable | 0.0 | 25.9 | 0.0 | 0.0 | 0.0 | 3.7 | 70.4 | 0.0 |
| shootable | 50.0 | 5.6 | 0.0 | 0.0 | 0.0 | 16.7 | 27.8 | 0.0 |

**Figure S4:** Confusion matrix showing the results for each of the three test sets obtained using the ResNet50 classifier. Rows show class labels with predicted class in columns.

| Test set 1 | null | clear | heavy-ppt | light-ppt | phase-sep | crystalline | optimisable | shootable |
|---|---|---|---|---|---|---|---|---|
| null | 94.6 | 0.3 | 2.0 | 2.0 | 0.3 | 0.3 | 0.0 | 0.3 |
| clear | 5.3 | 83.3 | 1.2 | 5.7 | 1.4 | 2.6 | 0.0 | 0.5 |
| heavy-ppt | 0.7 | 0.0 | 97.7 | 0.9 | 0.0 | 0.7 | 0.0 | 0.0 |
| light-ppt | 0.1 | 0.0 | 30.3 | 66.1 | 1.6 | 1.7 | 0.0 | 0.1 |
| phase-sep | 0.0 | 0.0 | 2.1 | 2.1 | 88.2 | 5.9 | 0.0 | 1.6 |
| crystalline | 0.0 | 0.3 | 1.3 | 2.4 | 2.4 | 91.3 | 1.1 | 1.3 |
| optimisable | 0.0 | 0.0 | 0.3 | 1.4 | 2.7 | 12.5 | 70.9 | 12.2 |
| shootable | 0.0 | 0.0 | 0.0 | 0.0 | 0.0 | 1.5 | 0.0 | 98.5 |

| Test set 2 | null | clear | heavy-ppt | light-ppt | phase-sep | crystalline | optimisable | shootable |
|---|---|---|---|---|---|---|---|---|
| null | 73.3 | 0.0 | 11.8 | 14.3 | 0.0 | 0.6 | 0.0 | 0.0 |
| clear | 4.9 | 71.6 | 4.2 | 12.9 | 2.5 | 3.0 | 0.0 | 1.0 |
| heavy-ppt | 0.9 | 0.0 | 94.3 | 2.0 | 0.0 | 2.4 | 0.2 | 0.2 |
| light-ppt | 0.2 | 0.2 | 26.5 | 67.5 | 1.4 | 4.0 | 0.1 | 0.2 |
| phase-sep | 1.0 | 0.0 | 4.4 | 2.4 | 81.0 | 9.5 | 0.7 | 1.0 |
| crystalline | 0.0 | 0.0 | 22.5 | 7.1 | 1.1 | 62.9 | 2.3 | 4.1 |
| optimisable | 0.7 | 0.0 | 18.5 | 3.0 | 5.2 | 15.6 | 31.1 | 25.9 |
| shootable | 0.0 | 0.0 | 0.0 | 0.9 | 0.0 | 0.9 | 0.0 | 98.1 |

| Test set 3 | null | clear | heavy-ppt | light-ppt | phase-sep | crystalline | optimisable | shootable |
|---|---|---|---|---|---|---|---|---|
| null | 25.0 | 0.0 | 50.0 | 16.7 | 8.3 | 0.0 | 0.0 | 0.0 |
| clear | 0.2 | 85.5 | 1.1 | 7.3 | 3.4 | 2.4 | 0.0 | 0.0 |
| heavy-ppt | 0.0 | 0.0 | 99.2 | 0.8 | 0.0 | 0.0 | 0.0 | 0.0 |
| light-ppt | 0.0 | 0.1 | 31.5 | 62.4 | 4.1 | 1.8 | 0.0 | 0.1 |
| phase-sep | 0.0 | 0.0 | 0.3 | 0.6 | 96.4 | 1.9 | 0.0 | 0.8 |
| crystalline | 0.0 | 0.0 | 0.0 | 4.4 | 5.5 | 89.0 | 1.1 | 0.0 |
| optimisable | 0.0 | 0.0 | 3.7 | 0.0 | 14.8 | 29.6 | 44.4 | 7.4 |
| shootable | 0.0 | 0.0 | 0.0 | 0.0 | 5.6 | 0.0 | 0.0 | 94.4 |

**Figure S5:** Confusion matrix showing the results for each of the three test sets obtained using the InceptionV3 classifier. Rows show class labels with predicted class in columns.

| Test set 1 | null | clear | heavy-ppt | light-ppt | phase-sep | crystalline | optimisable | shootable |
|---|---|---|---|---|---|---|---|---|
| null | 87.8 | 4.4 | 3.1 | 3.4 | 0.0 | 0.3 | 0.3 | 0.7 |
| clear | 1.2 | 88.5 | 1.2 | 6.5 | 0.2 | 2.2 | 0.2 | 0.0 |
| heavy-ppt | 0.2 | 0.0 | 97.2 | 1.8 | 0.0 | 0.7 | 0.0 | 0.0 |
| light-ppt | 0.0 | 0.0 | 24.6 | 72.2 | 0.2 | 3.0 | 0.0 | 0.0 |
| phase-sep | 0.5 | 0.0 | 3.7 | 1.1 | 79.7 | 12.3 | 0.5 | 2.1 |
| crystalline | 0.3 | 0.5 | 0.5 | 0.3 | 0.0 | 96.8 | 0.0 | 1.6 |
| optimisable | 0.3 | 0.3 | 0.7 | 0.7 | 0.0 | 14.5 | 75.3 | 8.1 |
| shootable | 0.8 | 0.0 | 0.0 | 0.0 | 0.0 | 0.0 | 0.0 | 99.2 |

| Test set 2 | null | clear | heavy-ppt | light-ppt | phase-sep | crystalline | optimisable | shootable |
|---|---|---|---|---|---|---|---|---|
| null | 72.1 | 1.9 | 8.7 | 11.8 | 0.0 | 3.7 | 0.6 | 1.2 |
| clear | 2.3 | 80.3 | 2.1 | 12.1 | 0.4 | 2.5 | 0.0 | 0.4 |
| heavy-ppt | 0.7 | 0.2 | 83.4 | 1.8 | 0.2 | 13.1 | 0.0 | 0.7 |
| light-ppt | 0.3 | 0.1 | 14.4 | 78.3 | 0.3 | 6.6 | 0.1 | 0.0 |
| phase-sep | 0.0 | 0.3 | 6.4 | 11.9 | 57.0 | 22.4 | 1.7 | 0.3 |
| crystalline | 0.0 | 0.4 | 3.4 | 3.8 | 0.0 | 90.6 | 0.4 | 1.5 |
| optimisable | 0.0 | 0.0 | 7.4 | 6.7 | 2.2 | 32.6 | 33.3 | 17.8 |
| shootable | 0.9 | 0.9 | 0.9 | 0.0 | 0.0 | 5.6 | 0.0 | 91.6 |

| Test set 3 | null | clear | heavy-ppt | light-ppt | phase-sep | crystalline | optimisable | shootable |
|---|---|---|---|---|---|---|---|---|
| null | 25.0 | 8.3 | 8.3 | 50.0 | 0.0 | 0.0 | 8.3 | 0.0 |
| clear | 0.0 | 90.4 | 0.2 | 7.8 | 1.0 | 0.5 | 0.2 | 0.0 |
| heavy-ppt | 0.2 | 0.0 | 98.3 | 1.2 | 0.0 | 0.3 | 0.0 | 0.0 |
| light-ppt | 0.0 | 0.0 | 25.8 | 70.7 | 1.2 | 2.4 | 0.0 | 0.0 |
| phase-sep | 0.0 | 2.5 | 1.6 | 4.7 | 84.1 | 7.1 | 0.0 | 0.0 |
| crystalline | 0.0 | 2.2 | 1.1 | 1.1 | 0.0 | 95.6 | 0.0 | 0.0 |
| optimisable | 0.0 | 11.1 | 7.4 | 0.0 | 7.4 | 18.5 | 55.6 | 0.0 |
| shootable | 0.0 | 0.0 | 0.0 | 0.0 | 0.0 | 0.0 | 0.0 | 100.0 |

**Figure S6:** Confusion matrix showing the results for each of the three test sets obtained using the Xception classifier. Rows show class labels with predicted class in columns.
